# Mercury contamination of an introduced generalist fish of intermediate trophic level

**DOI:** 10.1101/2022.12.01.518449

**Authors:** D. P. Gedig, M. Hauger, D. A. Armstrong, K. M. Jeffries

## Abstract

Mercury contamination is a global issue because mercury concentrations in aquatic systems are influenced by both natural and anthropogenic pathways, including the burning of fossil fuels and flooding during hydroelectric development. Mercury biomagnifies in aquatic ecosystems, leading to higher concentrations in piscivore fishes than those at lower trophic levels. Here, liver and muscle total mercury (THg) concentrations in black crappie *Pomoxis nigromaculatus* from three lakes in southeastern Manitoba, Canada were related to age, morphology and physiological traits to better understand the dynamics of mercury accumulation in an introduced generalist fish species. Black crappie liver and muscle samples from Big Whiteshell Lake (relatively large lake, 17.5 km^2^; n=30), Caddy Lake (small lake surrounded by wetlands, 3.1 km^2^; n=42) and Lac du Bonnet (river widening influenced by hydroelectric dams, 84.0 km^2^; n=29) were analyzed for THg content. These THg concentrations were then compared to black crappie mercury concentrations in other Canadian water bodies to assess within species relative contamination levels, as well as to mercury concentrations in other freshwater fishes to examine biomagnification. Age and size had strong positive correlations (r≥0.60) with muscle mercury concentrations. No evidence of acute point source contamination was found in the study area when compared to black crappie muscle mercury concentrations in other water bodies, and tissue THg concentration was not correlated with a reduction in gonadosomatic index (GSI) or hepatosomatic index (HSI). Analysis of liver THg in addition to muscle THg revealed the possible impacts of seasonal and ontogenetic differences in diet on exposure. Furthermore, THg analysis of liver and muscle tissue showed how generalist foraging techniques of black crappie may curb the progressively greater mercury exposure and resultant physiological consequences expected from ontogenetic diet shifts from invertebrates to fishes. Although there appeared to be temporally varied levels of mercury exposure (i.e., liver THg) by sex, there was no sex effect observed in long-term accumulation in the muscle. Flood risk is believed to be a key driver of differences in black crappie THg concentrations between lakes in the region. Black crappie bioaccumulated less mercury at age than primary piscivore species in the region. These results will help foster a better understanding of mercury biomagnification in boreal shield lakes within a region impacted by legacy mercury.

## Introduction

Mercury is a contaminant of global concern due to its dynamics within aquatic environments (Driscoll et al. 2013). While mercury is found naturally within ecosystems, it is also emitted by anthropogenic processes such as metallurgy, burning of fossil fuels and manufacturing (Pirrone et al. 2010). A primary form of mercury transport is via the atmosphere, and local and global trends in mercury emissions are reflected in ambient atmospheric concentrations even in remote areas (Wiklund et al. 2017). However, net deposition rates are dependent on various factors such as land cover and photoreduction, and natural processes including surface and groundwater flow, permafrost melt and wildfires are key in mobilizing and distributing mercury throughout the environment (Dastoor et al. 2022; Graydon et al. 2012; St. Louis et al. 2019). When inorganic mercury is transformed to methylmercury by anaerobic bacteria in aquatic conditions, mercury becomes more harmful to humans and other biota (Porvari & Verta 1995). Methylmercury is in an organic lipophilic state that facilitates bioaccumulation within an organism and biomagnification in the food web, resulting in tissue mercury levels that are higher than ambient concentrations (Borgmann & Whittle 1991; Lavoie et al. 2013). Because of bioaccumulation and biomagnification, consumption of fish is a major route of mercury intake for humans, where mercury has neurotoxic effects (Driscoll et al. 2013). Although the proportion of total mercury in fish that is methylmercury may vary with factors such as diet, age, and size, methylmercury is usually the main form found in adult fish muscle (Bloom 1992; Lescord et al. 2018). Methylmercury is also found in other tissues such as the liver, gonads and brain, despite generally having lower proportions of methylmercury to total mercury compared with proportions in muscle (Cizdziel et al. 2003; Khadra et al. 2019).

Mercury generally accumulates in freshwater fishes with increasing muscle mercury concentrations with increasing age and size (Phillips, Lenhart & Gregory 1980; Prestie et al. 2019; Sackett et al. 2013). Diet and trophic position are also key determining factors of mercury concentrations as piscivorous fish often accumulate mercury in their tissues faster than herbivorous fish due to biomagnification (Depew et al. 2013; Lavoie et al. 2013). Following mercury uptake in fish, the mercury is quickly eliminated from the liver, and then reallocated to the muscle where it has a longer retention time (Van Walleghem, Blanchfield & Hintelmann 2007). A comparison of the mercury concentration in both tissues helps understand long-versus short-term influences in exposure. Furthermore, this could provide insight into possible variation between lakes or by sex as has been observed in other freshwater fishes (Gewurtz, Bhavsar & Fletcher 2011; Madenjian et al. 2014; Nicoletto & Hendricks 1988). Fish can experience detrimental effects to their energetic and reproductive physiology due to exposure to mercury, which can manifest as reductions in the hepatosomatic index (HSI) and the gonadosomatic index (GSI), respectively (Friedmann et al. 1996; 2002; Larose et al. 2008). Examining these indices alongside tissue mercury levels allows an estimate of the metabolic impact of mercury on fish.

Black crappie *Pomoxis nigromaculatus* (Lesueur 1829) is a popular sportfish in North America that exhibits a generalist diet that ontogenetically transitions from one dominated by zooplankton into a piscivorous diet (Hanson & Qadri 1984; Keast 1968; 1985). The northernmost edge of the black crappie native range is found within southern Manitoba, Canada in the Red River and Winnipeg River drainages and within the south basin of Lake Winnipeg (Page & Burr 2011; Stewart & Watkinson 2004). Black crappie in Manitoba have a shorter spawning season that starts later in the year and have slower growth than more southern populations (Hauger 2020; Warren 2009). Black crappie have also been introduced into water bodies in Manitoba (Stewart & Watkinson 2004). This includes the notable population found in Whiteshell Provincial Park, Manitoba, which was thought to have been introduced in the 1940’s (D. Kroeker, *personal communications*). This black crappie population has since become more abundant and colonized the entire Whiteshell River system to the confluence with the Winnipeg River (Figure 1; D. Kroeker, *personal communications*). The lakes examined in the present study include Caddy Lake and Big Whiteshell Lake from within the Whiteshell River flowage, as well as Lac du Bonnet, which is a widening of the Winnipeg River (Figure 1).

**Figure 1.**
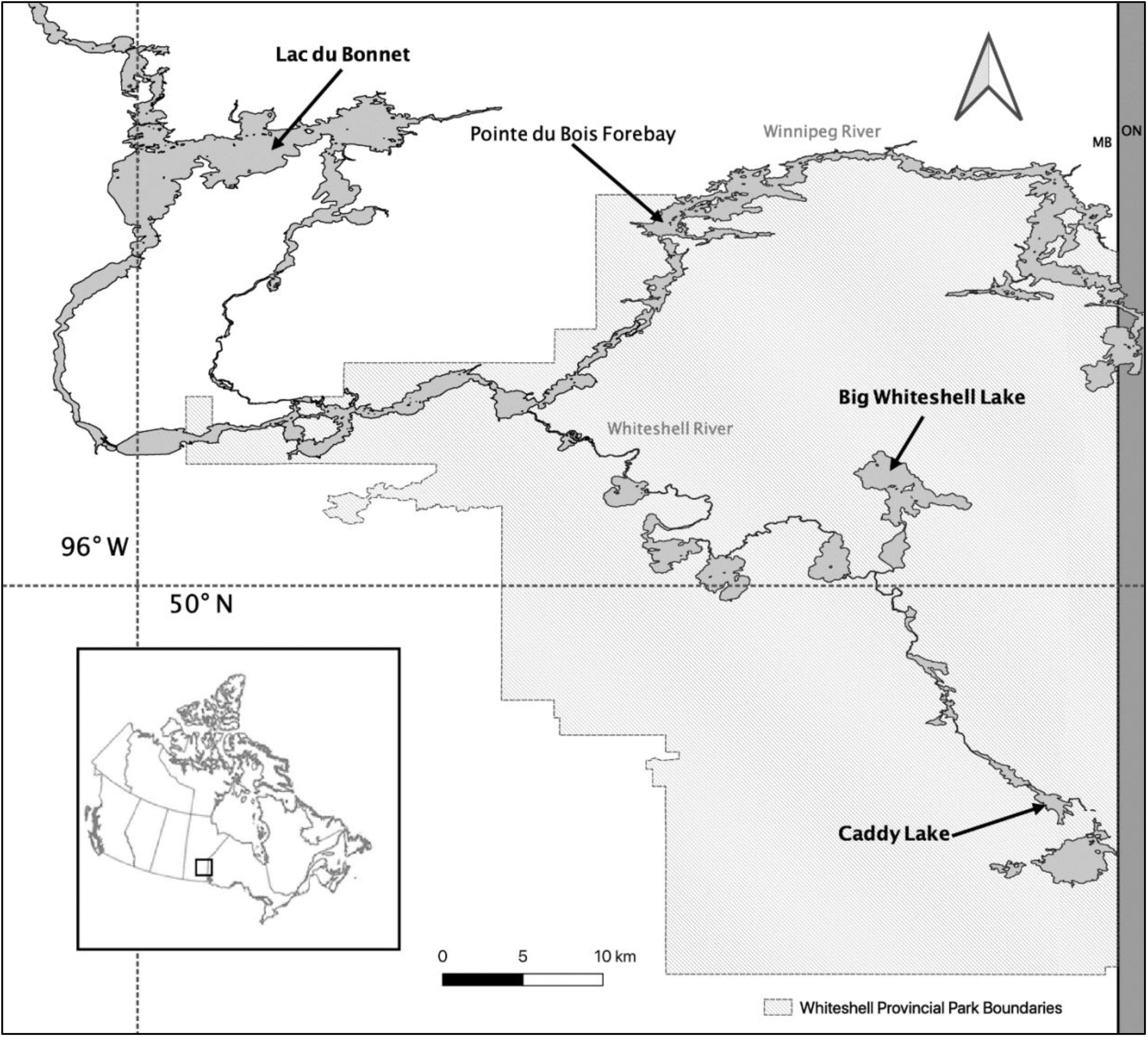
Map of relevant open waters in the Winnipeg River and Whiteshell River systems. Lake names in bold are those sampled from in this study. Inset map depicts Canada with a black box representing the area shown in detail. Shape files were sourced from the Manitoba Land Initiative (2001), and the map was created using QGIS (QGIS Development Team 2018).

Consideration of point sources of mercury and their potential for methylation is important for tracing mercury uptake. The lakes within Whiteshell Provincial Park are within a sub-basin of the Winnipeg River drainage, however as Lac du Bonnet is downstream of the Whiteshell River mouth, it has more potential external influences concerning land use and characteristics (Lake of the Woods Control Board 2000). From 1962 to 1969, a chlor-alkali plant in Dryden, Ontario released mercury effluent into the Wabigoon-English-Winnipeg River system, resulting in elevated mercury levels within biota downstream (Armstrong, Metner & Capel 1972). There is evidence that a legacy pool of mercury still contributes to elevated mercury concentrations in fish within the Winnipeg River, which were found to be higher than two other major tributaries of Lake Winnipeg (Jansen 2021). In addition, the Winnipeg River has several hydroelectric generating stations that alter its flow (Lake of the Woods Control Board 2000; Manitoba Hydro 2018). Rises in mercury inputs and the flooding that occurs from flow control are both linked to higher levels of mercury methylation, and thus mercury uptake in fishes (Orihel et al. 2007; Porvari & Verta 1995). Examining contemporary concentrations of black crappie will broaden the understanding of fish mercury levels in the Winnipeg River system and the lasting impact of mercury contamination.

This study investigated the dynamics of mercury accumulation in black crappie, an introduced fish species in Whiteshell Provincial Park that are of intermediate trophic level. Liver and muscle total mercury (THg) concentrations in black crappie from Caddy Lake, Big Whiteshell Lake and Lac du Bonnet were related to age, morphology and physiological traits to identify markers of mercury exposure and potential metabolic impacts. To this end, ontogenetic stage was quantified using age (assessed using otolith aging techniques) and measures of size (i.e., fork length and total mass) to determine the effect diet shifts have on mercury accumulation. We also examined whether there was a sex effect on mercury concentration. The black crappie data in this study were supplemented with those from Canadian government agencies to compare length-adjusted accumulation levels in various water bodies and characterize relative degrees of contamination within the species. Using additional data from the Coordinated Aquatic Monitoring Program (CAMP, campmb.com), the black crappie in this study were also compared to other freshwater fish species in the region to examine bioaccumulation by trophic level. We predicted that black crappie would have lower tissue mercury concentration than primary piscivore species due to differences in feeding behavior and a more generalist diet in black crappie. We also predicted that the black crappie from the three target lakes would not demonstrate acute contamination given the lack of prominent contemporary point sources in the study area, and that they would exhibit only minor, if any, physiological impacts due to mercury exposure as a result.

## Methods

### Study area

Of the three water bodies examined in this study, Caddy Lake is the smallest (3.1 km^2^ surface area), Big Whiteshell Lake is relatively large (17.5 km^2^ surface area) and the area of the Winnipeg River considered to be Lac du Bonnet is the largest (84.0 km^2^ surface area) (Pollom & Rose 2015; Wildlife and Fisheries Branch 2015a; 2015b). Whereas Lac du Bonnet is believed to be impacted by legacy mercury contamination, there are not any known major point sources of mercury to Caddy Lake and Big Whiteshell Lake (Jansen 2021). Caddy Lake has a higher risk of flooding than Big Whiteshell Lake due its smaller basin and restricted outflow (W. Kellas, *personal communications*). Caddy Lake also has greater coverage of wetlands in its watershed than Big Whiteshell Lake (Manitoba Land Initiative 2001). Although contemporary water quality data for the study area is limited, Big Whiteshell Lake appears to be more biologically productive than Caddy Lake (as measured by Secchi disk depth, total phosphorus concentration and chlorophyll *a* concentration; Water Quality Management Section 2020b).

### Sampling and morphological examination

Black crappie were terminally sampled from each of the three study lakes during 2019. Caddy Lake fish were collected using electrofishing (May 8^th^ and 14^th^), Big Whiteshell Lake fish were collected using trap nets (throughout August) and Lac du Bonnet fish were from gill nets (retrieved on September 17^th^). The sampling methods complied with the Canadian Council for Animal Care standards and guidelines as approved by the University of Manitoba Animal Care Committee (animal care permit F18-005 [AC11336]). Fork length (±1 mm) and total mass (±0.1 g) were measured and sex was determined by visual examination of the gonads for each fish. Only fish >140 mm in fork length possessing developed gonads were considered mature and assigned a sex. The mass of the liver and gonads (±0.1 g) was also recorded in fish >140 mm. Sagittal otoliths were extracted for aging and were reported in years as an integer. The otoliths were viewed whole against a black background under a dissecting microscope and each opaque band (annulus) was counted (Hauger 2020; Schramm & Doerzbacher 1982). Dorsal white muscle and liver tissue samples for mercury analysis were taken from each fish and stored at - 20ºC. Only fish ≤140 mm or >255 mm in fork length from Caddy Lake (n=42) and Big Whiteshell Lake (n=30) were included in this analysis. We assumed that there would be ontogenetic shifts in diet between the size classes and that the bigger group of fish would be at a reproductive age. The sample set from Lac du Bonnet (n=29) was not subject to this partitioning because we were limited to those caught in the gill net. Fork lengths ranged from 58 to 370 mm across the studied lakes.

### Mercury analysis

The total mercury (THg) concentration of the tissue samples was determined using cold vapor atomic absorption spectroscopy (CVAAS) with thermal decomposition and gold amalgamation. The instrument used was the Hydra II_C_ direct mercury analyzer by Teledyne Leeman Labs (Hudson, NH), which has a stated instrument detection limit of 0.001 ng (Teledyne Technologies Incorporated 2015). Calibration curves were constructed using the Certified Reference Materials (CRMs) MESS-4 and PACS-3, ranging from 0 ng (blank) to 400 ng of THg. The limit of quantitation (LOQ) was set as the lowest non-zero standard on the method calibration curve. Samples were first introduced into the decomposition furnace where they were dried (300ºC, 70 sec) and combusted (800ºC, 70 sec), then into the catalyst furnace (600ºC, 70 sec), and finally through the drying tube and gold amalgamation trap (600ºC, 30 sec) before entering the spectrometer. Samples were not homogenized nor digested before analysis. Once thawed, a portion of each tissue sample was placed in a nickel boat and weighed on an analytical balance (±0.0001 g). The mean amounts (± standard deviation) of muscle and liver tissue analyzed were 0.0591 ± 0.0104 g and 0.0546 ± 0.0150 g wet mass, respectively. Dissection tools and surfaces were cleaned with ethanol following handling of each sample. A concentration was calculated using the measured wet mass of the analyzed sample and the THg detected by the instrument. For THg measurements below the method LOQ, the concentration used in data analysis was calculated using 50% of the LOQ. Two muscle and four liver sample concentrations were calculated this way, which was considered acceptable as it was a small proportion (3.0%) of all reported mercury measurements (n=202).

For quality assurance, at least two CRMs and at least one duplicate measurement were performed for every ten tissue samples analyzed. The CRMs primarily used were MESS-4 (90 ± 40 ng g^-1^), DORM-4 (412 ± 36 ng g^-1^) and DOLT-5 (440 ± 180 ng g^-1^). Mean percentage recoveries (± standard deviation) were 105.3 ± 12.7%, 104.1 ± 2.4% and 99.5 ± 9.2%, respectively. Also, NIST 2709a (900 ± 200 ng g^-1^) was analyzed on a limited basis which served as a validation the calibration range. Duplicate measurements of the same sample needed to have a relative percent difference of 10% or be within 10 ng g^-1^ of their mean concentration to be considered satisfactory. If two quality assurance measurements were found unsatisfactory within the same set of ten samples, the instrument was assessed for recalibration and reanalysis was performed where necessary to validate measurements.

### Statistical analyses

The HSI and GSI were calculated to represent energetics and reproductive health, respectively (Equation 1; 2). Fulton’s Condition Factor (K) was calculated to represent the overall condition of a fish (Equation 3). The liver muscle index (LMI) was calculated as a ratio of liver THg concentration to muscle THg concentration (Equation 4). Data were grouped into two for analyses that were not mutually exclusive. The first included all fish from Big Whiteshell Lake and Caddy Lake to incorporate influences of ontogenetic shifts in diet on mercury dynamics between water bodies, hereafter referred to as the “ontogeny dataset.” The second included all mature fish >140 mm in fork length from all three lakes to investigate physiological impacts and the possible presence of a sex effect, hereafter referred to as the “adult dataset.” All statistical analyses were performed using RStudio (RStudio Team 2021).

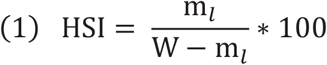

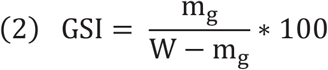

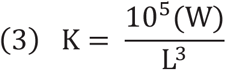

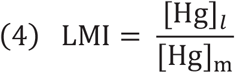

W = total wet mass (g)

L = fork length (mm)

m_*l*_ = liver wet mass (g)

m_*g*_ = gonads wet mass (g)

[Hg]_*l*_ = liver total mercury concentration (ng g^−1^)

[Hg]_m_ = muscle total mercury concentration (ng g^−1^)

A Pearson’s correlation matrix was constructed for each dataset to determine the strength and direction of variable associations. Grouped scatterplots with regression lines and residual plots were generated for each variable pair and were visually inspected for linearity and homogeneity of variance. Logarithmic transformations (base ten) were applied to one or both variables if needed to linearize data or address heteroscedasticity for an analysis of variance. The three lakes were expected to have differing biological and physicochemical conditions. To address this, all variables were centered by lake of origin following transformation by subtracting the group mean within each lake from each value (Equation 5). If after a transformation the data still did not meet the assumptions of a Pearson’s correlation, those correlations were omitted from the matrix and subsequent analyses.

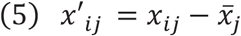

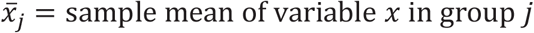

*x*_*ij*_ *= i*th value of variable *x* in group *j*

*x*′_*ij*_ *=* centered *i*th value of variable *x* in group *j*

Relationships with an absolute correlation coefficient of 0.5 or greater were fit with linear regression lines by sex and by lake of origin for the adult and ontogeny datasets, respectively. One-way Analysis of Covariance (ANCOVA) models were used to determine if the regression lines differed in slope or intercept. For the adult dataset, the factor of interest was sex, and tests were performed for each lake separately to account for varying conditions. The factor of interest was lake of origin for the ontogeny dataset, and thus tests were performed using the dataset as a whole. Type II sum of squares was used and the critical *P*-value was 0.05 after confirming that the slopes of the relationships did not statistically differ. To interpret the influence of differing slopes, covariate values where 95% confidence bands did not overlap were considered different. Outliers were not omitted from statistical analyses if instrumental error was not suspected in order to avoid bias and retain relationships as close as possible to those existing in nature. However, for the adult dataset, outlier values of liver THg concentration (>5 standard deviations from group-centered univariate mean) and the derived LMI from one fish were omitted from ANCOVA testing due to significant alteration of regression relationships. Pairwise deletion was used for any missing or omitted data. Where applicable, normality of residuals was investigated by visual inspection of histograms.

Measurements of muscle THg concentration in black crappie from three additional water bodies were acquired for comparison to those in this study. The Red River is part of the native range of this species, and although it does not show signs of acute point source mercury contamination, its drainage basin is subject to flooding which could promote mercury uptake (Brigham, Goldstein & Tornes 1998; Sando et al. 2003; Stewart & Watkinson 2004). Swan Lake is a lake within the Winnipeg River system upstream from Lac du Bonnet, which was exposed to the same mercury effluent during the 1960’s as Lac du Bonnet and is also affected by a proximate hydroelectric generating station (Lake of the Woods Control Board 2000; Tam & Armstrong 1972). Lake St. Clair was impacted by mercury effluent until the early 1970’s due to chemical plants located in Sarnia, Ontario, and sport fishes within the lake exhibited acute contamination thereafter (Gewurtz et al. 2010). To adjust for fork length, regression lines were fit for each water body with a logarithmic transformation of muscle THg concentration to satisfy linearity and homoscedasticity assumptions. The estimated value and 95% confidence interval were determined for 255 mm fish and used for comparison.

In addition, data on three other freshwater fishes from CAMP (2019) found in Manitoba, Canada were used to compare interspecific trends of mercury accumulation. Black crappie were sampled from Lac du Bonnet in this study, whereas northern pike *Esox lucius* L. 1758, walleye *Sander vitreus* (Mitchill 1818) and lake whitefish *Coregonus clupeaformis* (Mitchill 1818) were sampled from a comparable upstream location (Pointe du Bois Forebay). We examined muscle THg to age relationships because age is the preferred measure for comparing mercury bioaccumulation due to the confounding effects of differing length-at-age relationships between species (Neumann & Ward 1999). Linear regression lines were fit for each species with a logarithmic transformation of muscle THg concentration to satisfy linearity and homoscedasticity assumptions. The linear regression lines were then visually inspected to examine qualitative differences in mercury concentration at age.

## Results

### Adult dataset

Muscle THg concentration correlated with age (*P<*0.001, r=0.60), fork length (*P<*0.001, r=0.72) and total mass (*P<*0.001, r=0.70), but did not differ significantly by sex in any of the lakes. The full correlation matrix for the adult dataset is presented in Table 1.

**Table 1.** Correlation matrix for the adult dataset (n=74–80). Fish were sampled from Big Whiteshell Lake, Caddy Lake and Lac du Bonnet and have fork lengths >140 mm. Data was group-mean-centered by lake of origin. Pearson’s correlation coefficient (r) is reported in the top section of the matrix, and *P*-values are reported in the bottom section.

|  | 1 | 2 | 3 | 4 | 5 | 6 | 7 | 8 | 9 |
| --- | --- | --- | --- | --- | --- | --- | --- | --- | --- |
| **1. Age** | – | 0.76 | 0.79 | -0.05 | 0.00 | 0.15 | 0.60† | 0.28 | -0.24 |
| **2. Fork Length** | <0.001 | – | 0.94 | -0.24 | -0.11 | 0.08 | 0.72† | 0.15 | -0.38 |
| **3. Total Mass** | <0.001 | <0.001 | – | -0.03 | 0.04 | 0.17 | 0.70† | 0.12 | -0.39 |
| **4. K** | 0.66 | 0.03 | 0.81 | – | 0.23 | 0.32 | -0.07 | -0.16 | 0.09 |
| **5. HSI** | 0.99 | 0.36 | 0.75 | 0.05 | – | 0.64 | 0.01 | -0.04 | 0.03 |
| **6. GSI** | 0.19 | 0.48 | 0.14 | 0.005 | <0.001 | – | 0.18 | -0.12 | -0.10 |
| **7. Muscle [THg]** | <0.001† | <0.001† | <0.001† | 0.51 | 0.90 | 0.12 | – | 0.34 | -0.28 |
| **8. Liver [THg]** | 0.01 | 0.18 | 0.30 | 0.17 | 0.72 | 0.29 | 0.002 | – | 0.65 |
| **9. LMI** | 0.04 | 0.001 | <0.001 | 0.41 | 0.77 | 0.38 | 0.01 | <0.001 | – |

† log<sub>10</sub>(Muscle [THg]) data transformation was applied

Muscle THg concentration also correlated with liver THg concentration (*P=*0.002, r=0.34). Liver THg concentration correlated with age (*P*=0.01, r=0.28). Males had a greater intercept for LMI regressed on liver THg concentration in Big Whiteshell Lake (*P=*0.004) and Caddy Lake (*P=*0.04) than females (Figure 2). The LMI correlated with age (*P=*0.04, r=-0.24), fork length (*P=*0.001, r=-0.38), and total mass (*P<*0.001, r=-0.39).

**Figure 2.**
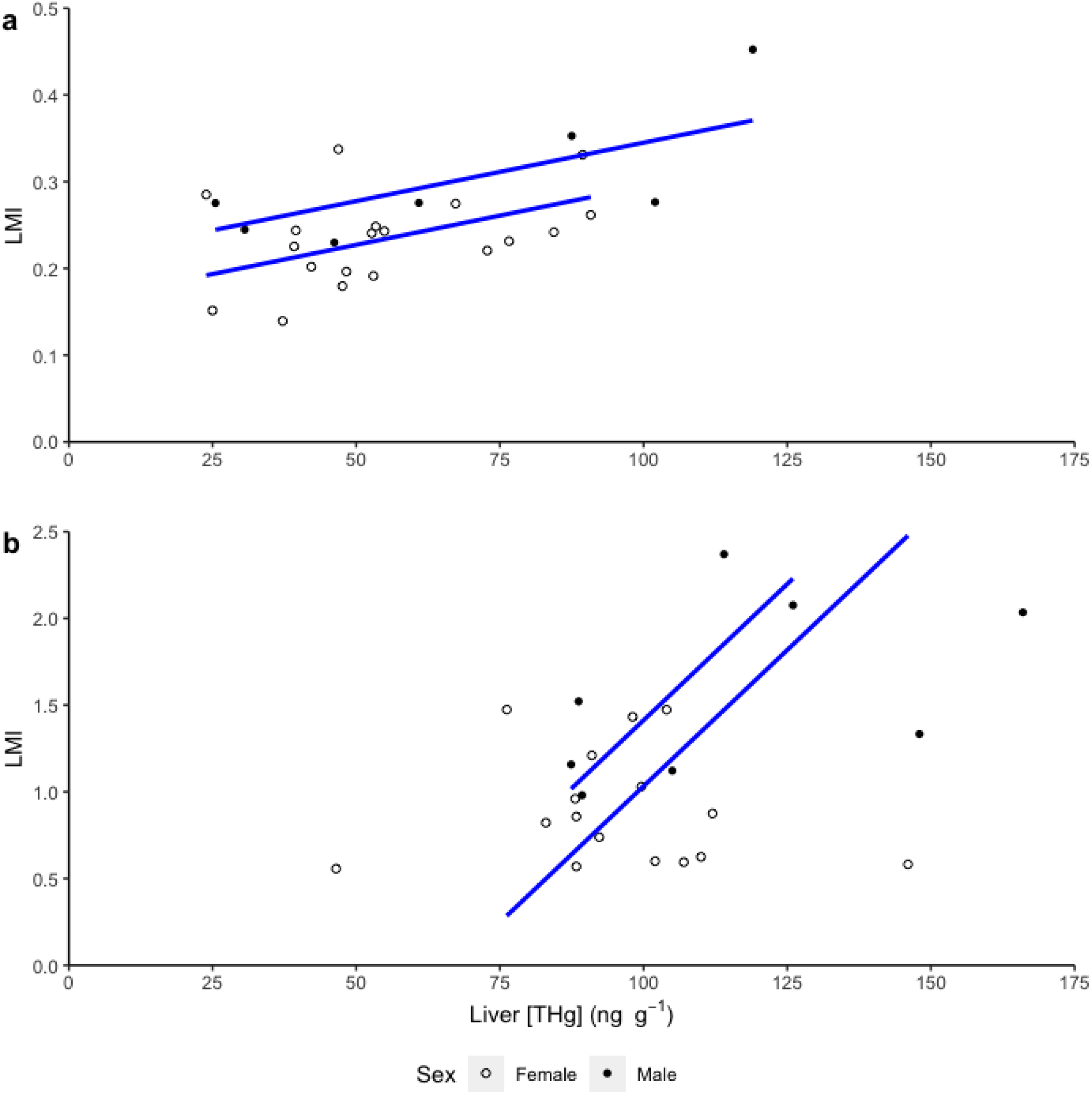
The LMI regressed on liver total mercury concentration in black crappie (*Pomoxis nigromaculatus*) sampled from Caddy Lake (a; n=26) and Big Whiteshell Lake (b; n=24). Fish were sampled in Manitoba, Canada and have fork lengths >255 mm.

### Ontogeny dataset

Muscle THg concentration correlated with age (*P*<0.001, r=0.83), fork length (*P*<0.001, r=0.93) and total mass (*P*<0.001, r=0.94). The full correlation matrix for the ontogeny dataset is presented in Table 2. Black crappie in Big Whiteshell Lake increased muscle THg concentration with age at a higher rate (*P<*0.001), but had lower THg concentration than fish from Caddy Lake until values converged at age eight (Figure 3). Muscle THg concentration increased with fork length (*P*<0.001) and total mass (*P*=0.02) faster in Big Whiteshell Lake, but black crappie in Caddy Lake had greater THg concentration values overall. Muscle THg concentration also correlated with liver THg concentration (*P*<0.001, r=0.76) and LMI (*P*<0.001, r=-0.82), and Big Whiteshell Lake had a greater intercept than Caddy Lake for both variables when regressed on muscle THg concentration (*P<*0.001 both). Age correlated with liver THg concentration (*P*<0.001, r=0.72) and LMI (*P*<0.001, r=-0.68) (Figure 3). Liver THg concentration increased at a higher rate with age in black crappie from Big Whiteshell Lake (*P*=0.02), and was greater than Caddy Lake after age three (Figure 3). The LMI decreased at a lower rate with age in Caddy Lake (*P*=0.002), but was greater overall in Big Whiteshell Lake (Figure 3). Liver THg concentration also correlated with K (*P*=0.02, r=0.28).

**Table 2.** Correlation matrix for the ontogeny dataset (n=72). Fish were sampled from Big Whiteshell Lake and Caddy Lake and have fork lengths <140 mm or >255 mm. Data was group-mean-centered by lake of origin. Pearson’s correlation coefficient (r) is reported in the top section of the matrix, and *P*-values are reported in the bottom section. Omitted relationships are indicated by “*NA*.” Table S1 presents applied data transformations.

|  | 1 | 2 | 3 | 4 | 5 | 6 | 7 |
| --- | --- | --- | --- | --- | --- | --- | --- |
| **1. Age** | – | NA | NA | NA | 0.83 | 0.72 | -0.68 |
| **2. Fork Length** | NA | – | 0.998 | 0.81 | 0.93 | NA | NA |
| **3. Total Mass** | NA | <0.001 | – | 0.84 | 0.94 | NA | NA |
| **4. K** | NA | <0.001 | <0.001 | – | NA | 0.28 | -0.08 |
| **5. Muscle [THg]** | <0.001 | <0.001 | <0.001 | NA | – | 0.76 | -0.82 |
| **6. Liver [THg]** | <0.001 | NA | NA | 0.02 | <0.001 | – | 0.23 |
| **7. LMI** | <0.001 | NA | NA | 0.49 | <0.001 | 0.05 | – |

**Figure 3.**
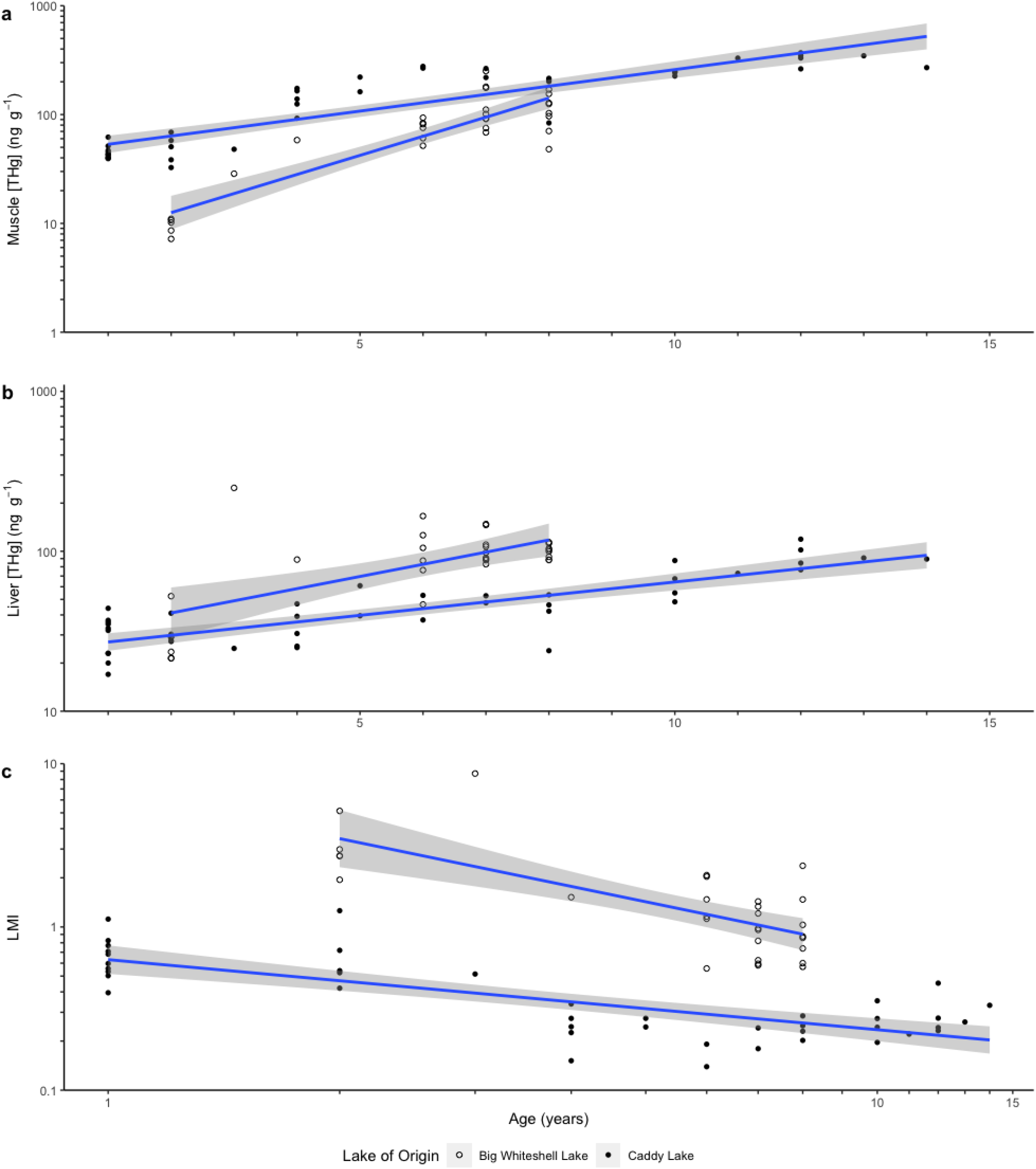
Muscle total mercury concentration (a), liver total mercury concentration (b) and LMI (c) regressed on age in black crappie (*Pomoxis nigromaculatus*). Fish were sampled from Big Whiteshell Lake (n=30) and Caddy Lake (n=42) in Manitoba, Canada and have fork lengths <140 mm or >255 mm. A 95% confidence band is shown for each linear regression line. Note the differing scale for plot C.

### Spatiotemporal and interspecific comparisons

A comparison of black crappie muscle mercury levels in three additional water bodies to those observed in this study is presented in Table 3. When adjusted to a length of 255 mm, Big Whiteshell Lake had the lowest mean mercury concentration and Lake St. Clair had the highest (Table 3). The confidence intervals for Caddy Lake, Swan Lake and the Red River overlapped, and the confidence interval of Lac du Bonnet overlapped with the Red River only (Table 3). The muscle mercury concentration of four freshwater fishes regressed on age showed that northern pike had the greatest mean mercury concentration at age, followed by walleye, then black crappie and then lake whitefish (Figure 4).

**Table 3.**
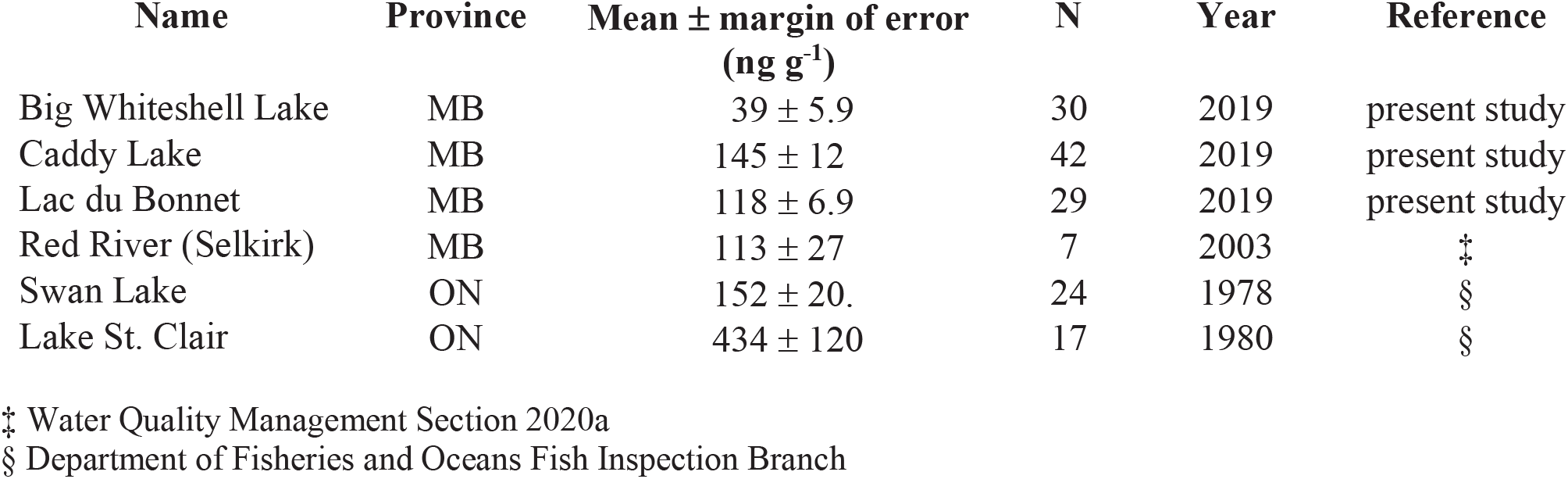
Black crappie muscle total mercury concentrations from various water bodies in Canada adjusted to a fork length of 255 mm. Margin of error is 95% confidence.

**Figure 4.**
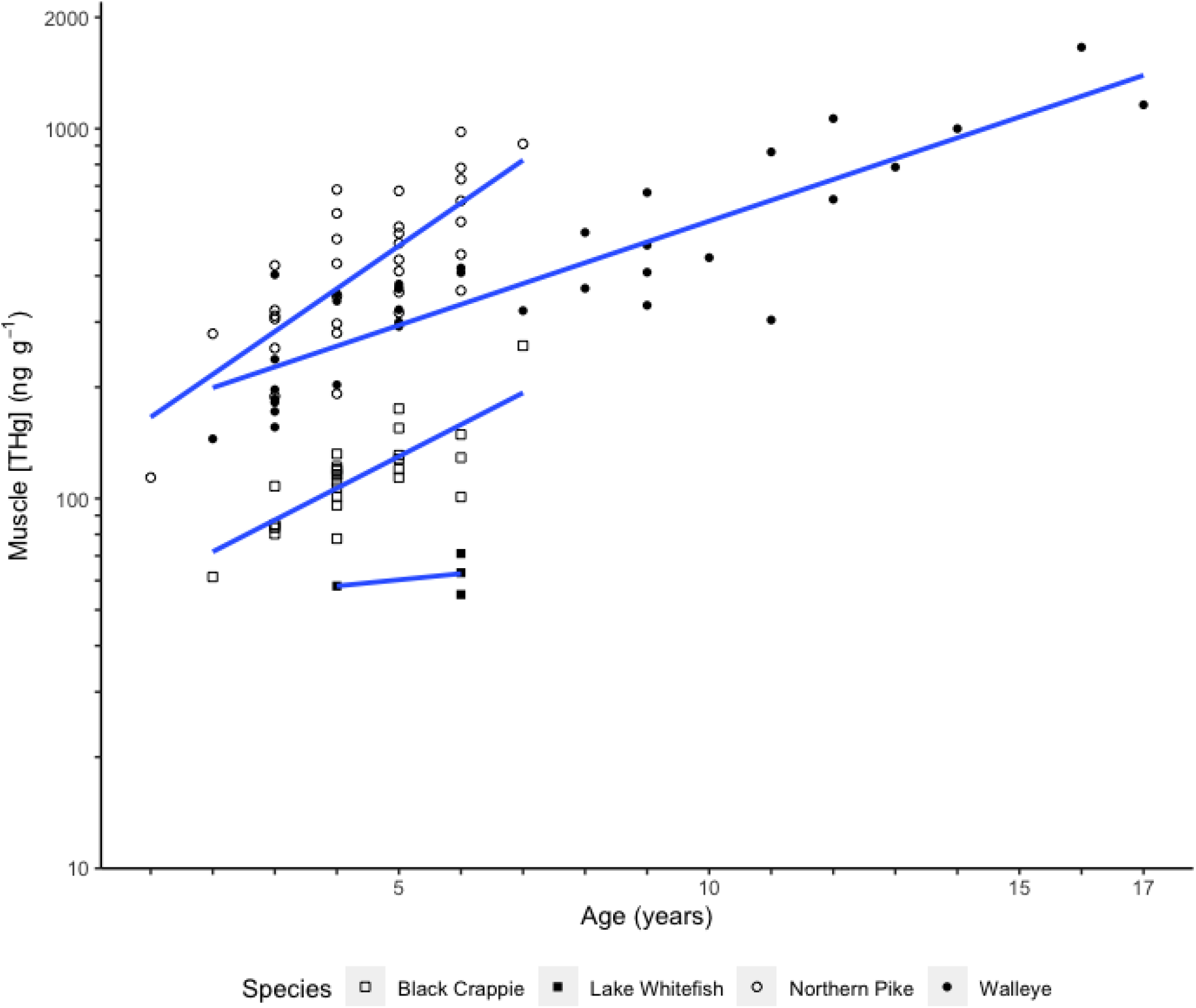
Muscle total mercury concentration regressed on age in four freshwater fish species from Manitoba, Canada. Black crappie (*Pomoxis nigromaculatus*) were sampled from Lac du Bonnet in 2019 (present study), and northern pike (*Esox lucius*), walleye (*Sander vitreus*) and lake whitefish (*Coregonus clupeaformis*) were sampled from the Pointe du Bois Forebay in 2016 (CAMP 2019). Linear regression lines are plotted for each species individually.

## Discussion

### Age, morphology and physiology

As black crappie and other piscivores increase in age and length, they opportunistically eat a diet consisting of larger prey and more fish (Keast 1985). Consumption of prey that is of higher trophic level results in increased muscle mercury concentrations (Depew et al. 2013; Phillips, Lenhart & Gregory 1980; Sackett et al. 2013). Similarly, the results of this study show increased mercury concentration in white muscle of black crappie with increases in age, length and mass. Following uptake, mercury is first distributed to the liver for detoxification, and any remaining mercury not initially eliminated (predominantly of the methylmercury form) is then quickly reallocated to muscle tissue for long-term storage (Burrows & Krenkel 1973; Cizdziel et al. 2003; Van Walleghem, Blanchfield & Hintelmann 2007). Liver mercury concentration could then reflect the degree of mercury exposure around the time of sampling, whereas muscle mercury concentration would represent the long-term effects of mercury retention and accumulation. Liver THg concentration increased with age in this study, likely due to the consumption of progressively larger prey with higher levels of mercury over time. Cizdziel et al. (2003) suggested such rises in liver mercury concentration with increased exposure are caused by greater demethylation rates. In contrast, liver THg concentration did not correlate with fork length or total mass in adult fish across the study area. Furthermore, while the LMI in freshwater fishes generally increases with higher trophic level, the LMI in black crappie had a negative association with age and growth (Cizdziel et al. 2003; Goldstein, Brigham & Stauffer 1996). Black crappie have an opportunistic generalist diet that is more dependent on prey availability than body size (i.e., gape) limitation, and it appears the fish sampled for this study did not shift food sources to manifest the rise in short-term mercury exposure observed in previous studies (Gaeta et al. 2018; Hanson & Qadri 1984; Keast 1968).

Elevated mercury concentrations are associated with physiological impairment in freshwater fish, including reductions in GSI and HSI (Friedmann et al. 1996; 2002; Larose et al. 2008). No association was found between either of these indices and tissue THg concentration, suggesting no deleterious impacts on fish physiology in the present study. Tissue mercury concentration has also been shown to negatively correlate with fish condition factor as a result of emaciation or hindered protein synthesis (Cizdziel et al. 2002; 2003; Suns & Hitchin 1990). The evidence of a positive association of body condition with tissue mercury concentration in this study is contrary to trends reported in other species. Provided sufficient prey abundance and modest environmental mercury, black crappie may overcome the metabolic cost of mercury exposure that can reduce fish condition, despite the continued elevation of tissue THg concentrations. A review of dose-response literature found minimal negative impacts at tissue levels of 200 ng g^-1^ (Dillon, Beckvar & Kern 2010). Whereas Big Whiteshell Lake fish exhibited muscle THg concentrations below this 200 ng g^-1^ threshold, both Lac du Bonnet and Caddy Lake had levels in excess, the latter having levels well above 300 ng g^-1^. However, there were still no negative effects of mercury accumulation on the HSI, GSI and condition observed in this study.

### Sex effect

Muscle THg concentrations did not significantly differ between males and females of similar age and morphology, which suggests that the accumulation of mercury in black crappie is not influenced by sex overall. However, males in Big Whiteshell Lake and Caddy Lake had a significantly greater LMI value than females for any given level of liver contamination. In black crappie, the gonads have ripened and are ready for reproduction by late autumn, a strategy permitting an early spring spawn after minimal winter foraging (Bevelhimer & Breck 2009; Suski & Ridgway 2009). At the same time, HSI increases (perhaps in preparation for reproduction and overwintering) and mesenteric fat stores are depleted (Beuchel, Marschall & Aday 2013; Hauger 2020). Males begin increasing their gonad sizes in August, which is earlier than females (Beuchel, Marschall & Aday 2013). If males initially invest greater energy into developing the gonads than the liver, females could have a relatively greater increase in HSI. Reduction in liver size (e.g., due to reduced condition) has been correlated with increased liver mercury concentrations as a result of the slow elimination rate of mercury (Cizdziel 2003). Vice versa, if female black crappie have an increase in HSI that is not found in males at the time of sampling, it may explain the relatively greater liver mercury levels of males in Big Whiteshell Lake sampled in August. Whereas females invest more energy in gonadal development overall, males provide parental care of spawning which, due to limited feeding opportunities, requires survival on stored energy almost exclusively (Bevelhimer & Breck 2009; Cooke et al. 2006). This stress could also result in relative reduction in HSI for males, which could account for the marginally significant greater liver mercury levels in males of Caddy Lake during May. Sexually dimorphic detoxification, such as unequal elimination rates as found by Madenjian et al. (2014), could also impact these disparities, however it was not measured in this study. Nevertheless, any differences in behavior or physiology between sexes appear to ultimately balance out, resulting in similar rates of mercury accumulation in muscle tissue in the long term.

### Spatiotemporal and interspecific comparison

In 1971, higher mercury contamination was detected in Caddy Lake fish (dorsal muscle concentrations higher than 500 ng g^-1^) compared with Big Whiteshell Lake (Tam and Armstrong 1972). A similar pattern was observed in this study, where muscle THg concentrations were higher in Caddy Lake versus fork length and total mass. Caddy Lake has a higher risk of flooding than Big Whiteshell Lake, and when soils are flooded, the resultant anaerobic conditions favor bacterial mercury methylation and subsequently promote uptake in aquatic food webs (Porvari & Verta 1995). Furthermore, there is greater coverage of wetlands in the Caddy Lake watershed, which has also been linked to increased levels of mercury methylation (Hurley et al. 1995; Rypel 2010; Selvendiran et al. 2008). Big Whiteshell Lake appears to be more biologically productive than Caddy Lake, which is associated with lower food web mercury concentrations via dilution across greater biomass quantities (Lescord et al. 2019; Rypel 2010). Black crappie in Big Whiteshell Lake have a higher observed growth rate than Caddy Lake, which is also linked to lower fish mercury levels (Borgmann & Whittle 1991; Hauger 2020). It appears that the differences in muscle mercury concentrations between Caddy Lake and Big Whiteshell Lake are due to differing limnological conditions rather than point source contamination. The disparity seems to decrease with age as muscle mercury levels in both lakes converge at age eight.

Liver mercury levels appeared to follow a contrary pattern, with fish from Big Whiteshell Lake having higher concentrations than Caddy Lake fish after age three. This could be due to seasonal variations in diet as black crappie will consume more fish and exhibit higher growth in late summer than in spring, which were the sampling periods of Big Whiteshell Lake and Caddy Lake, respectively (Haines 1980; Keast 1968). Younger fish are less likely to consume the higher trophic level prey that would cause an elevation of liver mercury levels, which may explain why the disparity between Caddy Lake and Big Whiteshell Lake appears after age three. In turn, the comparatively low muscle mercury levels coupled with the high liver mercury levels in Big Whiteshell Lake raised LMI values above those found in Caddy Lake. This may also indicate relatively greater consumption of young or small-bodied fishes in Big Whiteshell Lake, as these fishes exhibit lower proportions of methylmercury in muscle (Lescord et al. 2018). Unfortunately, the confounding effects of differing sampling periods prevents drawing conclusions about contrasting detoxification or exposure within the two lakes. However, they suggest the potential impacts of seasonal changes in diet for adult fish.

When comparing black crappie mercury levels in this study to other Canadian water bodies, Lake St. Clair muscle mercury concentrations were notably higher than the others due to acute point source contamination. Beyond this, Caddy Lake, Lac du Bonnet, Swan Lake and the Red River appeared to exhibit similar muscle mercury concentrations, despite having varying degrees of point source contamination. This similarity in mercury levels suggests that flood risk and the construction of dams are major factors affecting black crappie mercury levels in Manitoba. In addition, the comparable contamination present in native black crappie from the Red River suggests that the introduced population of Lac du Bonnet and Whiteshell Provincial Park exhibits similar mercury dynamics to native populations, despite the legacy pool of mercury within the region. The greater level of bioaccumulation with higher trophic level demonstrates the nature of biomagnification and the impact trophic interactions have on contamination. Historical study of mercury levels in the Winnipeg River system also indicated a disparity between lower trophic level fish, such as lake whitefish, and primary piscivores in levels of contamination (Derksen 1979).

### Limitations

The aim of this study was to assess mercury accumulation patterns in southeastern Manitoba. A larger sample size was prioritized to serve this goal, which was prohibitive of higher accuracy mercury analysis techniques (e.g., sample homogenization to reduce measurement variability). This sampling regime resulted in a relatively coarse numeration of tissue mercury concentrations which broadly represented the study area. In addition, fish sample selection was limited to catch success, which was not consistent between all three lakes and led to sampling occurring in each lake at different times throughout the open water season. Different sampling methods were used in each lake as each approach is best suited for certain times of year, but size-selectivity bias was introduced as a consequence (Hauger 2020). Despite using size-based sample partitioning and attempting to control for covariates to mitigate these factors, the strength of some observed patterns may have improved with a more uniform method of sample collection.

## Conclusions

Black crappie mercury levels in this study were consistent with established relationships of bioaccumulation, exhibiting muscle mercury concentrations that increased with growth and age. Analysis of liver mercury levels facilitated an understanding of the influence of seasonal and ontogenetic shifts in the black crappie diet in boreal lakes, the former being more influential in adults. It appeared that the black crappie in this study did not consume prey of a high enough trophic level to cause the increase in mercury exposure expected in fishes that shift food sources, which may also explain why there was no strong effect on HSI or GSI and no reduction in condition. Black crappie appear to be less vulnerable to contamination than fish species of higher trophic level, namely the piscivore specialist northern pike. Further, flood risk may be a primary factor that determined the level of contamination based on spatiotemporal comparison. The black crappie in this study were not adversely affected by their levels of mercury contamination based on the endpoints we measured. Nonetheless, given the unique trophic placement of black crappie as generalist feeders, understanding the dynamics of mercury within this species provides insight into how this contaminant impacts aquatic food webs.

## Contributions

D.P.G. and K.M.J. conceived the study idea. D.P.G. and M.H. sampled and gathered data. M.H. tabulated morphological and physiological data. D.P.G. performed mercury analysis with the aid of D.A.A. D.P.G. performed data analysis and prepared the manuscript with draft review by K.M.J.

## Acknowledgements

We would like to thank Michaela McKennitt for her efforts during our field season, as well as Dr. Feiyue Wang for permitting the usage of lab resources and for providing valuable comments. We are also grateful for the guidance given by Wolfgang Jansen of North/South Consultants Inc. Daniel Rheault and Andrew Burton of the Department of Agriculture and Resource Management (MB), Jennifer Van de Vooren of Manitoba Hydro (CAMP) and Dr. David Depew of Environment and Climate Change Canada provided much appreciated help in gathering supplemental datasets. This project was funded by a Manitoba Fish and Wildlife Enhancement Fund (FES #17-016) and a Natural Sciences and Engineering Research Council of Canada Discovery Grant (#05479) awarded to K.M.J.

