## Supporting Information for "Mercury contamination of an introduced generalist fish of intermediate trophic level"

**Table S1.** Logarithmic data transformations applied to variables for relationships in the Pearson's correlation matrix of the ontogeny dataset. Omitted relationships are indicated by "*NA*."

| | **1** | **2** | **3** | **4** | **5** | **6** | **7** |
|---|---|---|---|---|---|---|---|
| **1. Age** | | | | | | | |
| **2. Fork Length** | *NA* | | | | | | |
| **3. Total Mass** | *NA* | $\log_{10}(2,3)$ | | | | | |
| **4. K** | *NA* | $\log_{10}(2)$ | $\log_{10}(3)$ | | | | |
| **5. Muscle [THg]** | $\log_{10}(5)$ | $\log_{10}(5)$ | $\log_{10}(5)$ | *NA* | | | |
| **6. Liver [THg]** | $\log_{10}(6)$ | *NA* | *NA* | – | $\log_{10}(5,6)$ | | |
| **7. LMI** | $\log_{10}(1,7)$ | *NA* | *NA* | – | $\log_{10}(5,7)$ | – | |

**Table S2.** XLS file containing the data used in this study.
